# Neural Representations Behind ‘Social Norm’ Inferences In Humans

**DOI:** 10.1101/230508

**Authors:** Felipe Pegado, Michelle H.A. Hendriks, Steffie Amelynck, Nicky Daniels, Jessica Bulthé, Haemy Lee Masson, Bart Boets, Hans Op de Beeck

**Affiliations:** Department of Brain and Cognition, KU Leuven, 3000 Leuven, Belgium; Center for Developmental Psychiatry, Department of Neurosciences, KU Leuven, 3000 Leuven, Belgium; Leuven Autism Research consortium, KU Leuven, 3000 Leuven, Belgium

**Keywords:** multisensory, multivoxel pattern analysis, social appropriateness, social reasoning, mentalizing

## Abstract

Humans are highly skilled in social reasoning, e.g., inferring thoughts of others. This mentalizing ability systematically recruits brain regions such as Temporo-Parietal Junction (TPJ), Precuneus (PC) and medial Prefrontal Cortex (mPFC). Further, posterior mPFC is associated with allocentric mentalizing and conflict monitoring while anterior mPFC is associated with self-related mentalizing. Here we extend this work to how we reason not just about what one person thinks but about the abstract shared social norm. We apply functional magnetic resonance imaging to investigate neural representations while participants judge the social congruency between emotional auditory in relation to visual scenes according to how ‘most people’ would perceive it. Behaviorally, judging according to a social norm increased the similarity of response patterns among participants. Multivoxel pattern analysis revealed that social congruency information was not represented in visual and auditory areas, but was clear in most parts of the mentalizing network: TPJ, PC and posterior (but not anterior) mPFC. Furthermore, interindividual variability in anterior mPFC representations was inversely related to the behavioral ability to adjust to the social norm. Our results suggest that social norm inferencing is associated with a distributed and partially individually specific representation of social congruency in the mentalizing network.

## 1. Introduction

Humans have an extraordinary capacity to understand their conspecifics. This ‘social reading’ in natural environments involves the processing of visual cues – e.g., face expressions^1^, auditory cues – e.g., prosody ^2,3^, and other sensory information that is usable to infer others’ feelings, desires and thoughts ^4–5^. Although our mentalizing capacity (or Theory of Mind) typically relies on sensory cues of concrete targets, it can also be performed with more abstract cues such as verbal information about a person^7^. Mentalizing tasks systematically activate the so-called mentalizing brain network, including the temporoparietal junction (TPJ), precuneus (PC) and medial prefrontal cortex (mPFC) ^4–6,8^. However it is unclear how the human brain mentalizes at a more abstract level, for instance, when targeting not only the thinking of one particular person but instead how the general population ‘thinks’? In other words, how does the human brain infer what ‘most people’ think, for instance concerning appropriate social behavior?

From a behavioral point of view, it is now known that the development of such abstract inferences of social norms relies on active learning during concrete social interactions at very early ages (at least 3 years-old)^9^ (but see also^10^ for a passive social learning alternative). Learning social norms in a particular culture and in a particular family ultimately generates personal references of what most people think about appropriate reactions in different contexts (personal bias for social norms)^5,11,12^. Imagine for a moment that you are presenting your holiday pictures to an audience of relatives. Upon displaying a photograph of a beautiful scene, one observer reacts by expressing admiration using a vocal utterance (such as “uaaaauu”), whereas another expresses disgust (“uuuurg”). In this situation, the appropriateness of these two reactions will probably be perceived as congruent and incongruent, respectively, by most of us. Note however that in more nuanced or ambiguous situations, it can be challenging to judge social congruency and to further estimate the ‘common sense’ or the ‘social norm’ (i.e., what most people would think about the social congruency), as it can be difficult to be detached from our own perspective.

Here we present a new behavioral and neuroimaging paradigm which implements this latest example, requesting people to infer how most people would judge the congruency of vocal reactions to visual scenes. Here we will focus on the neural representations underlying this ‘social norm’ inferencing, an unexplored aspect in the social cognitive neuroscience literature.

We expect to find social congruency information represented in the core mentalizing network but not in sensory areas in the visual or auditory systems. In addition, the mPFC might show further dissociations, as self-related mentalizing (egocentric) has been associated with brain activity in more *anterior* parts of mPFC, while mentalizing about others (allocentric) has been related to activity in more *posterior* parts of mPFC^7,12,13^, as confirmed in a meta-analysis of more than 100 studies^14^. Furthermore, conflict monitoring is also hosted in posterior parts of mPFC^15^. Thus, monitoring conflict of social congruency itself and/or between self-related and allocentric social norm responses, could also engage the posterior part of mPFC. We therefore predict to find stronger social congruency representations in posterior mPFC than in anterior mPFC.

## 2. Results

### 2.1 Behavioral results during the fMRI

During the experiment in the fMRI scanner, binary judgments of social congruency (i.e., congruent vs incongruent) relative to the inferred social norm were collected. For each participant and run, behavioral similarity matrices were created by pairwise comparing responses for the 96 Audio-Visual conditions (12 visual X 8 auditory; see Fig.1A). The most common response across runs was calculated at the individual level (Fig. 1B) and at the group level (Fig. 1A). The group level result reflects the ‘shared social norm’ pattern of response among participants and follows essentially (in 94 out of 96 conditions) a cross-modal valence congruency pattern, i.e., visual and auditory stimuli with the same valence (both positive or both negative) are considered congruent while different valences are considered incongruent. Further, the correlation of these similarity matrices across 144 recordings (24 subjects X 6 runs; one subject excluded for missing run) revealed higher within-subject correlations (Pearson’s r =.54) than between-subject correlations (r = 0.29) (paired t-test: t(24) = 10.66; p <.0001). This result suggests that despite the fact that subjects were explicitly instructed to imagine ‘what most people would answer’, they were still more consistent with themselves than with others (personal bias). Importantly, there was a high degree of variability in the between-subject correlations, demonstrating that some participants were better than others in the ability to match the ‘shared social norm’ (see Figs. 1B and 3B). We will later use this between-subject variability in task performance to investigate interindividual differences in brain representations (see section 2.5).

**Figure 1:**
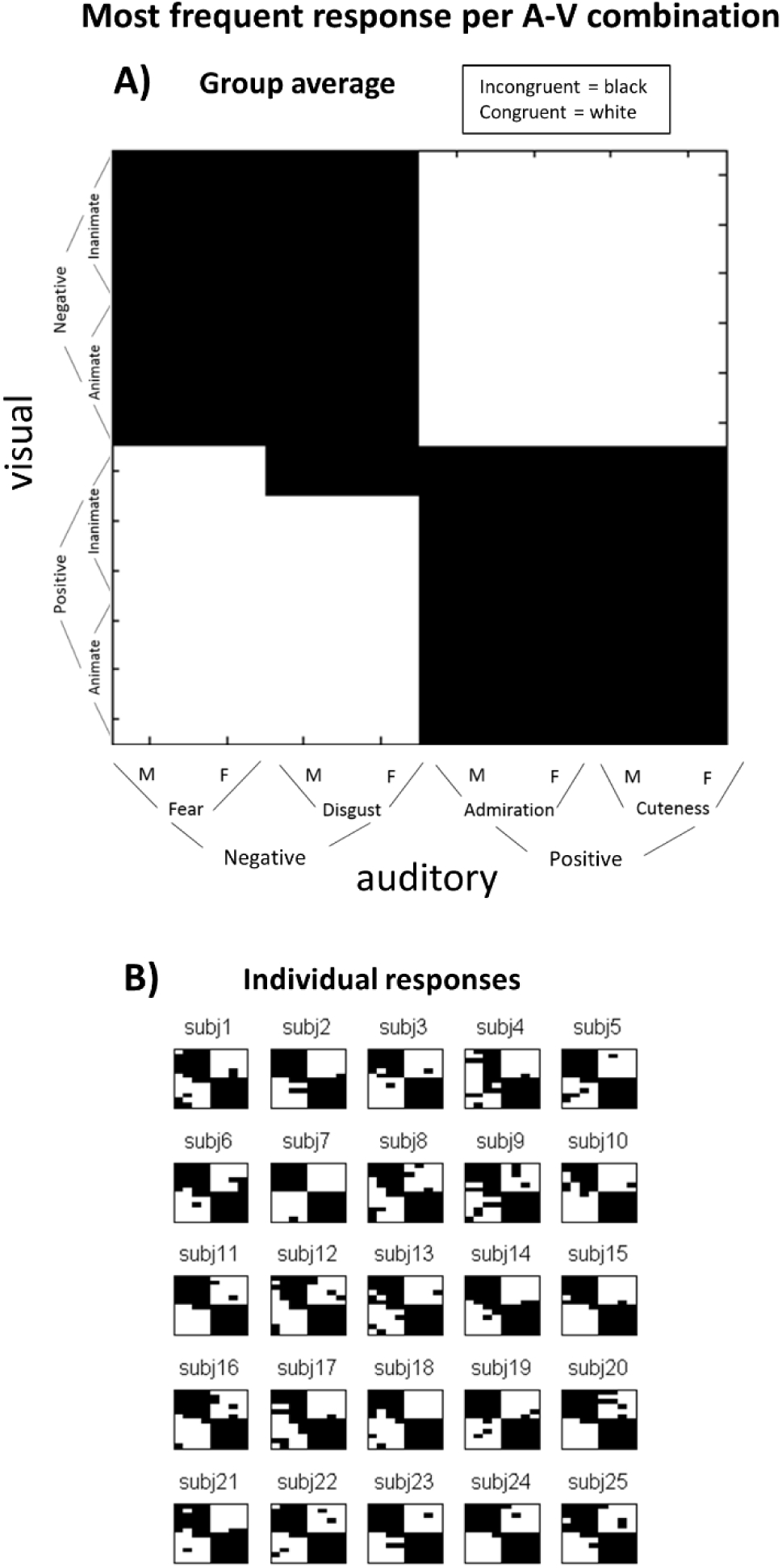
Behavioral responses during the scanner. Most frequent binary (congruent vs incongruent) response per Audio-Visual (A-V) stimuli combination (8 audio X 12 visual = 96) across the six runs, at the group level (A), representing thus the ‘shared social norm’ among participants, and at the individual level (B). Both for visual and auditory domains, half of the stimuli presented a positive valence and the other half a negative valence.

### 2.2 Behavioral results using a fine-grained scale (outside scanner)

Outside the scanner, 2 runs of the same task were administered, but instead of binary responses (congruent vs incongruent), subjects used a finer-grained 9-level scale to rate the congruency level (see Methods). The same pattern of results was found, i.e., higher within-subject correlations than between-subjects correlation (r =.67 vs.50 respectively; two-tailed paired t-test: t(24)= 6.67; p <.0001), again with variations between subjects in how much they correlate with other subjects in terms of which congruency responses they give to specific visual-auditory stimulus combinations. Thus, both using finegrained scales and binary decisions the pattern of result is the same.

### 2.3 Behavioral validation of the social norm perspective relative to egocentric perspective

We assumed here that between-subject correlations are a quantitative measure of the extent that individual participants can identify the shared social norm, which they were required to do through explicit task instructions. We tested this assumption by asking an independent group of participants to perform the same task with the fine-grained rating scale but now judging according to their own perspective (egocentric perspective) (see Methods). Results revealed that *within*-subject correlations were equivalent to those in the social norm perspective task (r =.65 vs.67 respectively; t(43) = −0.3) (see Fig. 2). More importantly however, egocentric perspective responses showed significantly lower *between*-subject correlations than in the social norm perspective (r =.33 vs.50; t(43) −7.1; p<.0001). In other words, by comparing the two fine-grained tasks, we find an objective increase in the similarity of the response patterns among participants when they are requested to judge social congruency under the ‘social norm’ perspective relative to when they use their personal perspective. This result demonstrates the sensitivity of our paradigm for subjective changes in the adopted perspective.

**Figure 2:**
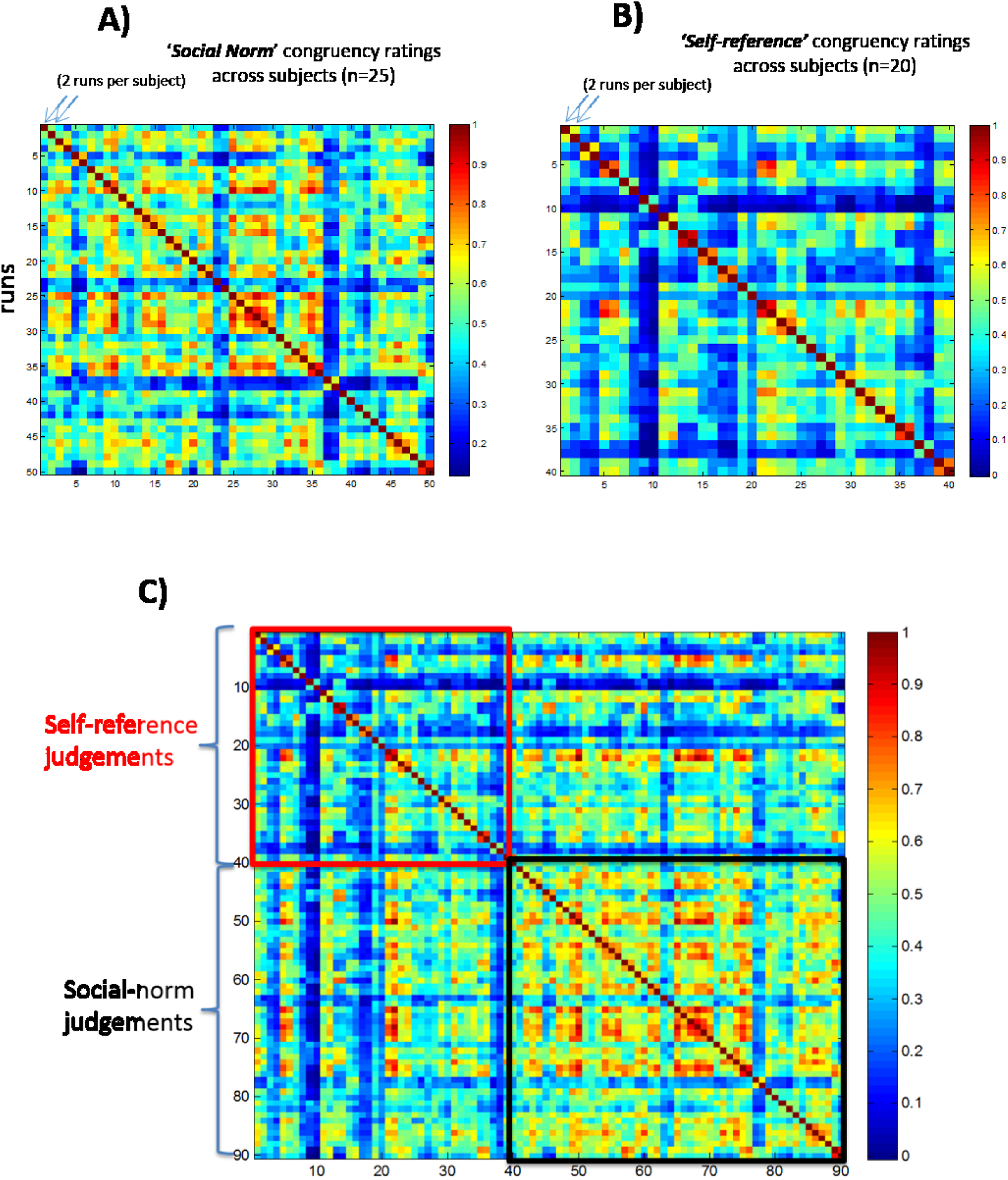
Behavioral responses outside the scanner using a fine-grained scale. A) *Same task as in the scanner (social norm reference)*. Outside the scanner, subjects performed 2 runs of the same social norm perspective task but instead of binary responses, they used here a more fine-grained scale (9 levels). B) *Control task (self-reference)*. A separate group of subjects performed a control task, where the judgements were based on their own perspective of social congruency, using again a 9 level scale (see Methods for details). C) *Comparing self-reference versus ‘social norm’ reference judgements*. Comparison of the results of the two tasks, so that each of the in total 45 participants is correlated with each of the other participants.

### 2.4 Neural representations of social norm processing

We used correlational MVPA to investigate whether there is a systematic difference between the multi-voxel pattern associated with congruent trials and the multi-voxel pattern associated with incongruent trials (see Methods), thus revealing a neural representation of (in)congruency with the social norm. We calculated the correlation between multi-voxel patterns elicited by the same condition, r(same) (e.g., correlating congruent with congruent), and compared it with the correlation between patterns from different conditions, r(diff) (e.g., correlating congruent with incongruent). As expected and shown in Figure 3A, most brain regions of the mentalizing network represent social congruency as revealed by r(same) being higher than r(diff): PC (p =.046; all p’s Bonferroni-corrected for multiple comparisons, i.e., the number of ROIs), TPJ (p =.01) and posterior mPFC (p =.009) (see Fig. 3A). However, this was not the case for the anterior mPFC (p >.9) nor for any of the sensory regions: EVC (p >.9), LOC (p >.9), EAC (p >.9), and TVA (p =.12).

**Figure 3:**
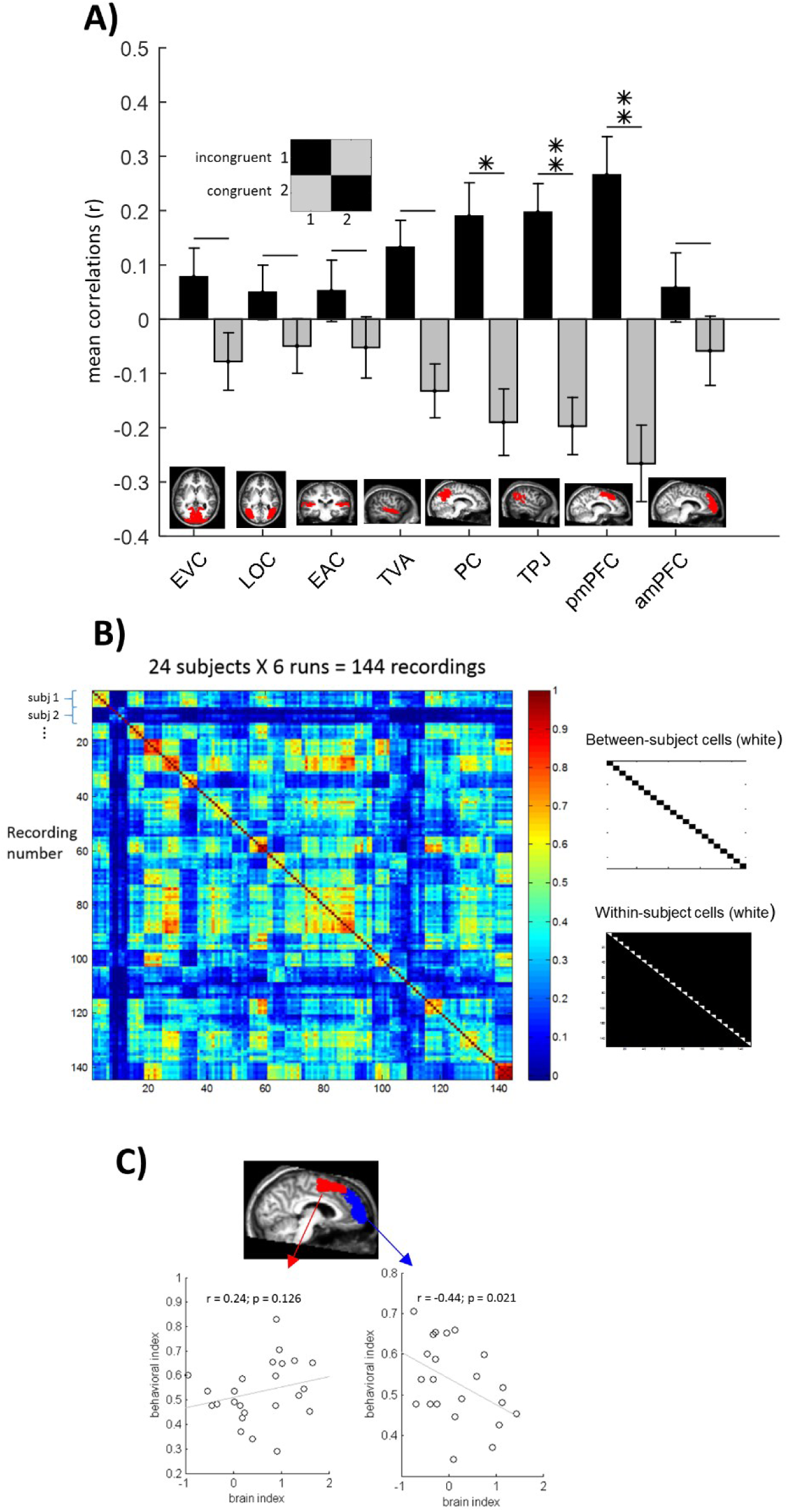
Social Congruency Brain Representations. Two conditions (congruent vs incongruent) were defined in a GLM model of the fMRI data and then 2 × 2 neural similarity matrices were created (inset). A) *ROIs with social congruency information*. None of the sensory areas (EVC = Early Visual Cortex; LOC = Lateral Occipital Complex; EAC = Early Auditory Cortex; TVA = Temporal Voice Area) show significant social congruency information content, while three of the mentalizing network ROIs did show (PC = Precuneus; TPJ = Temporo-Parietal Junction and mPFC = medial Prefrontal Cortex in its posterior part). B) *Behavioral response patterns across subjects*. Similarity of behavioral responses for the 96 audio-visual combination conditions, across runs and subjects (left panel) were calculated for each pair of subjects (Pearsons’ r correlations). Within and between subject correlations (cells in white; right panel) revealed higher within than between-subjects correlations, and an important variability across subjects. A behavioral indication of the ability to infer what most people would answer instead of oneself (a social norm mentalizing ‘performance’), was indexed by the individual between/within correlations ratio. C) *Linking neural and behavioral data*. To test if the social norm mentalizing performance (behavioral index) could be explained by differences in social congruency neural data in subparts of mPFC, brain x behavior correlations were performed.

### 2.5 Correlation between Neural x Behavioral data

The dissociation between posterior and anterior mPFC is directly relevant with regard to the perspective taken by our participants during the social congruency task. Previous studies have related allocentric perspective-taking with activity in posterior mPFC, and egocentric perspective taken with activity in anterior mPFC ^7,12–14^ We asked subjects to take an allocentric perspective, which might be the explanation for the presence of the neural representation in posterior mPFC and not in anterior mPFC. However, not all subjects performed this task in the same manner. As mentioned before, the size of between-subject correlations in behavioral response patterns, which we have shown to be modulated by the adoption of a social norm vs egocentric perspective (see Section 2.3), showed a lot of inter-subject variability. Hence, we wondered whether participants who were less able to adopt the requested allocentric social norm perspective may show a stronger (egocentric) congruence representation in anterior mPFC? To test this hypothesis, we computed a behavioral index quantifying to what extent the response pattern of a particular subject is similar to the other participants relative to him/herself: a between/within subject correlations ratio. A lower ratio indicates that a participant was less able to incorporate the shared social norm response. Results indeed revealed the predicted direction of the association: a negative correlation between social congruency information in *anterior* mPFC and the behavioral index of social norm inference: Pearson’s r =-.44; one-tailed p =.021 (Fig. 3C). In contrast, *posterior* mPFC shows a nonsignificant trend in the opposite direction (r =.24; p =.126). Comparing the differences in correlation between anterior and posterior mPFC, we found a significant difference (Z = −2.89; p = 0.002).

## 3. Discussion

To shed light upon the neural basis of ‘social norm’ inferences, we used a naturalistic audio-visual fMRI paradigm that mimics social reactions (vocalizations) in different visual contexts while asking subjects to imagine what most people would answer concerning the social congruency.

Behavioral data enabled us to (1) assess to what extent our paradigm is sensitive to the adopted perspective, with reference to an allocentric “social norm” versus an egocentric perspective, and (2) quantify the degree to which individual participants adhere to the shared social norm. First, behavioral response patterns revealed that when using the social norm perspective, participants objectively show higher between-subject correlations than when an egocentric perspective was adopted (Fig. 2),. Further, as there is no unequivocal ‘correct response’ on these subjective judgments of social congruency, similarity of response patterns across subjects was used to quantify task ‘performance’, i.e., the ability of each participant to match to the shared social norm (Fig. 1), revealing a good degree of variability across subjects (Fig. 3B). Importantly, *within*-subject correlations were higher than between-subject correlations in both social norm and self-related tasks and presented the same correlation level across the two tasks (Fig. 2).

At the brain level, we found that social congruency processing was not hosted in low and high-level visual and auditory sensory areas. In contrast, three of the core regions of the so-called mentalizing network ^4–6,8^ were engaged: Precuneus (PC), Temporo Parietal Junction (TPJ) and medial Prefrontal Cortex (posterior but not anterior part). Our allocentric mentalizing task could have favored the posterior instead of the anterior mPFC site for social information processing. Accordingly, previous work have shown a dissociation between egocentric and allocentric mentalizing processing in anterior and posterior mPFC respectively ^7,12–14^ Taken together, the recruitment of the core mentalizing network and the dissociation in mPFC suggests that largely the same neural mechanisms are used for mentalizing about a concrete single person and for more abstract general population mentalizing.

We further tested if the social norm perspective used during scanning would be related to the lack of social congruency information in anterior mPFC. If so, we would expect that participants who are less able to adopt an allocentric social norm perspective are the ones who show the strongest evidence for egocentric social congruency representations in anterior mPFC. And indeed, this is exactly the pattern that was observed in anterior mPFC (Fig. 3C). The pattern was clearly different in posterior mPFC.

We should acknowledge that the mere observation of a social congruency representation does not pinpoint which processes are reflected in this representation. For the same reason we cannot be sure to what extent the significant congruency representation in the three mentalizing regions reflects similar or different functions. As one example of potentially different functions, the posterior part of mPFC has also been associated with conflict monitoring ^15^, including conflict monitoring in the context of social information processing ^16–18^. Assuming that conflict monitoring would have played a role in posterior mPFC activity, it remains unclear whether posterior mPFC may have tracked conflicts concerning the social congruency itself (congruent vs incongruent) or rather conflicts between self-related and social norm responses (or even both). Further investigations should clarify this issue.

Taken together, the present results are compatible with the following interpretation: when inferring how most people would judge the appropriateness of social behavior, one engages the mentalizing network in the same way as when mentalizing about specific persons. During social norm mentalizing, three of the core mentalizing regions show a significant representation of social congruency. The fourth region, anterior mPFC, shows considerable inter-subject variability in this congruence representation which is related inversely to the ability to take the allocentric perspective.

## 4. Material and Methods

### 4.1 Participants

Twenty-five healthy subjects (7 female, 23.16 ± 3.32 years old, 7 left-handed) took part in the fMRI study. They all reported normal or corrected-to-normal vision, normal hearing and no neurological or psychiatric disorders. They received a financial compensation for their participation. The study was approved by the Medical Ethics Committee UZ/KU Leuven University and all methods were performed in accordance with the relevant guidelines and regulations. All participants provided written informed consent prior to scanning.

### 4.2 Visual stimuli

Twelve images were selected from the standardized and widely used emotional pictures set IAPS (International Affective Picture System) ^21^, based on extreme valence ratings (positive vs negative) and on animacy categorization (animate vs inanimate). The six animate (e.g., humans, animals…) and the six non-animate pictures (e.g., landscapes, objects…) were orthogonal to the image valence, with half of them rated positively (e.g., happy baby), and half of them negatively (e.g., people being threatened with a gun). Based upon the quality of the brain responses evoked by a larger dataset of 24 IAPS images present in the pilot fMRI study (cf. infra), this final set of 12 images was selected: numbered 2341, 1710, 1750, 5760, 5825, 7492, 3530, 1300, 1930, 9290, 9300, 9301 in the IAPS database. IAPS policy requests not to publish the original images.

### 4.3 Auditory stimuli

Eight different non-verbal vocal utterances were used, inspired by previous work^22^. They express four different emotional reactions that could be more or less congruent with the pictures previously selected. Utterances expressing disgust, fear, admiration and cuteness, were recorded in an expressive but still natural manner (not an exaggerated caricature). Each emotional vocalization was performed by one male and one female actor. They were recorded in a sound-proof room at 96 kHz sampling rate and 32-bit resolution, and were down-sampled to 44 kHz and 16-bit mono-recordings to reduce the size of the audio files. All stimuli had a fixed duration of 700 ms and an equivalent total Root Mean Square (RMS) power (−17.4 +/− 0.17 dB). Stimuli were slightly manipulated in Cool Edit Pro software and Adobe Audition CC 2015 software. Identical 600 ms silent periods were added before the onset of each auditory stimulus to create a natural delay from the visual stimulus onset, and a 100 ms silent period was added after the end of the utterance to provide stable ending transitions. Stimuli can be found here: https://osf.io/t7xp9/?viewonly=74aa08cefd634f09a70bac531ecf880e.

### 4.4 Behavioral Task in the scanner

Participants were lying in the scanner while watching a visual display and hearing auditory input through headphones. They were instructed to imagine they were seeing images (photographs) together with other unknown people. For each image, after a short delay, participants heard a vocal reaction (emotional utterance) that could be more or less congruent with the particular scene. The task was to evaluate the congruency of the vocal reaction in relation to the visual context. Yet, they did not have to judge this congruency from their own personal perspective, but instead they were explicitly instructed to evaluate whether most people would consider this vocal response appropriate or not (mentalizing the ‘social norm’) and respond accordingly. The two assigned buttons (congruent vs incongruent) were switched after 3 runs out of 6, and the assigned order was balanced across subjects.

### 4.5 Experimental fMRI runs

The fMRI session consisted of 6 runs, each with 96 pseudo-randomly presented experimental trials, i.e., all 12 visual stimuli paired with all 8 auditory stimuli. Additionally, 10 silent fixation trials were included among them, as well as 3 initial and 3 final dummy trials, making a total of 112 trials, with 4.5 seconds of Stimulus Onset Asynchrony (SOA), summing to 504 seconds of duration per run. Each experimental trial started with a visual image for 2.5 secs during which an auditory utterance was played via headphones (from 0.6 to 1.3 secs relative to the onset of the visual image) to simulate a natural delay before the vocal reaction. A 2 secs fixation cross was then displayed until the end of the trial. Subjects could respond any time within the trial and were instructed to press the buttons as soon as they know the answer.

### 4.6 fMRI data acquisition

Imaging data were acquired using a 3T Philips Ingenia CX scanner (Department of Radiology of the University of Leuven) with a 32-channel head coil. Each functional run consisted of T2*-weighted echoplanar images (EPIs), with voxel size = 2.52 × 2.58 × 2.5, interslice gap 0.2 mm, TR = 2550 ms, TE = 30 ms, matrix = 84×82, 45 slices, field of view (FOV) = 211 × 211 × 121. In addition to the functional images we collected a high-resolution T1-weighted anatomical scan for each participant (182 slices, voxel size = 0.98 × 0.98 × 1.2 mm, TR = 9.6 ms, TE = 4.6 ms, 256 × 256 acquisition matrix). Stimuli were presented using Psychtoolbox 3 (Brainard, 1997). Visual stimuli were displayed via an NEC projector with a NP21LP lamp that projected the image on a screen the participant viewed through a mirror. Viewing distance was approximately 64 cm. Auditory stimuli were presented through headphones at a comfortable hearing level.

### 4.7 fMRI preprocessing

Imaging data were preprocessed and analyzed using the Statistical Parametrical Mapping software package (SPM 8, Welcome Department of Cognitive Neurology, London, UK) and MATLAB. Functional images underwent slice timing correction (ascending order; first image as reference), motion correction (3rd degree spline interpolation), co-registration (anatomical to functional images; mean functional image as reference), and spatial normalization to the standard MNI (Montreal Neurological Institute) brain space. Functional images were resampled to a voxel size of 2.2 × 2.2 × 2.7 mm and spatially smoothed by convolution of a Gaussian kernel of 5 mm full-width at half-maximum^23^. One run of one subject was not considered due to excessive head movement (cut-off: 1 voxel size for two successive images).

### 4.8 General Linear Model (GLM)

We applied a general linear model focusing upon the representation of social congruence. For each participant and run, pre-processed images were modeled for each voxel using GLMs. They included regressors for each experimental condition and the 6 motion correction parameters (x, y, z for translation and rotation). Each predictor’s time course was convolved with the canonical hemodynamic response function (HRF) in SPM. The social congruency GLM had two conditions (congruent vs incongruent) based on cross-modal valence congruency (e.g., a positive image combined with a negative utterance is considered ‘incongruent’). This approach guaranteed a perfect balance between ‘congruent’ and ‘incongruent’ conditions, while avoiding potential visual or auditory biases, because all visual and auditory stimuli occurs an equal number of times for congruent and incongruent trials. As we used extreme ends of the valence continuum (very positive or very negative), both for the visual and auditory domains, we assumed that in binary choices this social congruency criterion would reflect quite well the participants’ responses. Indeed, group average responses showed a large overlap with this a priori definition of social congruency, as it was the most common response in 94 out of 96 A-V combinations (see Fig. 1A). Thus, this social congruency modeling also represents to a very large extent, the ‘shared social norm’ pattern, among the participant sample. The two social congruency conditions were modeled in relation to the time-window during which social congruency judgements could be performed, i.e., from the beginning of the auditory presentation (0.6 secs from trial onset) until the end of the trial (4.5 secs).

### 4.9 Regions of Interest (ROIs)

As primary ROIs we targeted the mentalizing neural network: medial Prefrontal Cortex (mPFC), Temporo-Parietal Junction (TPJ) and Precuneus (PC). Following the same approach as a recent meta-analysis of different mentalizing tasks^6^, we used the same parcels that were obtained in functional connectivity studies, both for mPFC^28^ and TPJ^29^. Note that only right hemisphere parcels are available for these two ROIs. Further, as we did not have particular hypotheses for the two subdivisions of TPJ (anterior and posterior parcels) we grouped them together in a single ROI. In contrast, for the mPFC, we kept this distinction as the literature shows a clear functional dissociation between anterior vs posterior parts for self-related vs others-related mentalizing processes respectively ^7,12–14^ by integrating the four original parcels into two. Finally, for PC, we used the anatomical mask in WFU Pickatlas (SPM).

In addition, we included two visual and two auditory ROIs as further control ROIs. Our approach was to select the best available templates to delineate brain areas corresponding to low-level as well as high-level processing in each modality, avoiding in that way both manual delineation of ROIs and the use of several functional localizers. Low-level processing is localized in primary sensory cortices, of which we know that anatomical landmarks provide a proper approximation. Thus, we used anatomical masks from the anatomical atlas WFU PickAtlas Toolbox (Wake Forrest University PickAtlas, http://fmri.wfubmc.edu/cms/software). The low level visual ROI (Early Visual Cortex-EVC) was defined based on Brodman’s areas (BA) 17 and 18 as they are widely accepted landmarks for low level visual processing. The low-level auditory ROI (Early Auditory Cortex-EAC) was composed by BA 41 and 42. The resulting EVC and EAC ROIs presented very thin configurations. This would lead to unrealistic delimitations of early processing cortex given the spatial uncertainty involved when comparing brains across subjects. We thus made them thicker by 1 voxel in all three directions to accommodate the spatial uncertainty in the probabilistic map. This procedure is nowadays already incorporated in PickAtlas through the 3D dilatation function.

Pure anatomical delimitation is less appropriate for high-level sensory regions, thus we used functional parcels obtained independently by other laboratories. As a high-level visual ROI, a functional parcel of the Lateral Occipital Complex (LOC) from the Kanwisher lab was used ^24^. The high-level auditory ROI was based upon the ‘Temporal Voice Area’- TVA probabilistic map from Belin’s lab ^25^ concerning more than 200 subjects, available at neurovault (http://neurovault.org/collections/33/).

These general masks were combined (by means of a conjunction analysis) with individual functional data that specify voxels modulated by our task: the F-contrast of all task trials against fixation trials, at a threshold of 0.0001 (uncorrected for multiple comparisons), using a separate ‘neutral GLM’ where all task trials were modeled as a single condition (fixation was implicitly modeled). ROIs with at least 20 active voxels were included. If a given participant ROI did not meet these criteria, his/her data was not used in the group analysis for this ROI. This situation only took place for two subjects in the anterior mPFC.

To ensure that no overlap occurs between ROIs, we visually inspected ROI borders of each ROI pair and restricted the ROIs to avoid the overlap. As a first measure, we restricted the TVA probabilistic map to the most significant voxels by imposing an arbitrary threshold of t = 50, which restricted the ROI to their classical temporal cortex disposition, and reduced considerably its overlap with other regions (e.g., TPJ). The resulting map was then transformed in a binary mask. As an additional measure, for this and all the other ROIs (all binary masks), we excluded the remaining overlapping voxels from the largest ROI of each pair. Only the following ROI intersections presented some overlap: EVC × LOC, EVC × PC, EAC × TVA, EAC × TPJ and TVA × TPJ (the first of each pair being the largest one). This procedure ensured a complete separation of the ROIs.

Given its role in emotional processing, we have also considered using an amygdala ROI. Yet, we are not confident that our standard imaging protocol at 3T was sensitive to pick up multi-voxel patterns in amygdala, which was further confirmed by null results in other (here unreported) analyses focusing upon other dimensions such as the properties of the visual/auditory stimuli. The ROI did not show any significant representation of social congruence, but given our doubts about the data quality in this ROI we do not think we can interpret this result and we preferred to not include the ROI further.

### 4.10 Correlation-based multivoxel pattern analysis

We used correlation-based multivoxel pattern analysis (MVPA) to explore how the spatial response pattern in individual ROIs differs between experimental conditions^30^. For each participant, we extracted the parameter estimates (betas) for each condition (relative to baseline) in each voxel of the ROIs. These obtained values for each run were then normalized by subtracting the mean response across all conditions (for each voxel and run separately), to emphasize the relative contribution of each condition beyond global activation level, as previously done in the literature^31,32^. The full dataset (6 runs) were randomly divided into two independent subsets of runs (using ‘randperm’ function in matlab). Thus, typically three runs were randomly assigned to set 1 and three other runs to the set 2 of the classification procedure. In the single case of incomplete data (5 runs instead of 6), only two runs were assigned as set 1. The multivoxel patterns associated with each condition (congruent and incongruent) in set 1 (runs averaged) were pairwisely correlated with the activity patterns in set 2 (runs averaged) by using the ‘corrcoefʼ function in matlab (Persons’ r correlation coefficient). This procedure of splitting the data in two parts followed by correlating the multi-voxel patterns was repeated 100 times. The final 2×2 neural similarity matrix for each ROI was obtained by averaging these 100 matrices.

To test whether a certain region contained information about social congruency, we applied the following procedure. First, we calculated for each ROI, the mean correlations in the diagonal (correlation of the same condition across runs) and non-diagonal cells (correlation of different conditions across runs) of the neural similarity matrix. Then, we performed a two sample two-tailed t-test across participants for diagonal vs non-diagonal mean correlations. This procedure is based on the fact that the same condition will typically show higher similarity across runs relative to different conditions, i.e., higher correlations for diagonal vs non-diagonal cells^34^. Lastly, a Bonferroni correction for multiple comparisons (i.e., the number of ROIs) was applied.

To investigate the potential relationship between brain information level of social congruency with behavioral ‘performance’ on the social norm inference task, i.e., the ability to match the response chosen by their peers, we performed a correlational analysis using a brain index (diagonal minus non-diagonal values) against a behavioral index (individual between/within subjects correlation ratio; see Fig. 3A).

### 4.11 Searchlight MVPA analysis

Searchlight MVPA analysis was used as a complementary way to check for potential missing anatomical areas outside our a priori ROIs. We used the searchlight scripts of the cosmo MVPA toolbox^35^ to search for local neighborhoods that revealed significant congruence representations (diagonal > nondiagonal), using the default parameters (e.g., spherical neighborhood of 100 voxels). The searchlight analysis was performed for each individual participant, after which the results were smoothed to 8 mm full-width at half-maximum and a 2^nd^ level model was performed (both using SPM). One participant was excluded from this analysis because of a missing run (excluded for excessive movement). No extra brain region captured the social congruency information.

### 4.12 Univariate analysis

Correlations between activity patterns (MVPA) are not necessarily related to overall differences in activity level between conditions (univariate analysis). Nevertheless, it is relevant to know whether MVPA findings are found in the context of effects that can also be picked up by a univariate voxel-level difference in activity (FWE corrected at p <.05). There were no significant effects of social congruency (congruent trials higher or lower than incongruent trials) in a whole brain analysis.

### 4.13 Outside the scanner

Prior to scanning, participants performed valence rating judgments for the stimuli of each modality (visual and audio) separately, using a 9-level scale. This was used to familiarize participants with the stimuli set used in the fMRI runs and to create group averaged perceived valence models.

Additionally, participants also performed two runs of the main task (as inside the scanning, i.e., using an allocentric ‘social norm’ perspective to judge congruency) but instead of binary responses (congruent vs incongruent) they used here a 9-level scale rating. One run was performed before and one after the fMRI scans. Beyond the familiarization with the task, this was helpful to check reproducibility of response patterns within and between-subjects, in a more fine-grained way.

### 4.14 A behavioral control experiment: Allocentric perspective

To validate the sensitivity of our behavioral paradigm to the ‘social norm’ perspective taken during the main behavioral and neuroimaging experiments, we compare these results with a behavioral control experiment. The control experiment included the same task but instead of asking participants to judge ‘what most people would answer’ (allocentric perspective), they were instructed to answer as a function of their own perspective (egocentric perspective). Twenty other subjects (age: 19.9 +/− 2.02 yrs old) performed this task with a 9 level rating of congruence. The visual stimulus set was larger (double) but the analysis presented here was restricted to the exact same stimulus set used in the main fMRI experiment and in the task performed outside the scanner previously described. The auditory set was perfectly equivalent at the perceptual level, but slightly differed in terms of low-level auditory features such as duration and total RMS.

### 4.15 Data Availability

The files needed to replicate the analyses will be made available on the Open Science Framework (e.g., ROI definitions and individual subject representational similarity matrices). Other aspects of the data (raw data files, other steps in the analyses) are available from the corresponding author on reasonable request.

## Acknowledgments

Stefania Bracci and Jean Steyaert for helpful discussions, Anne de Vries and Ninke De Schutter for their help in behavioral data acquisition and The Department of Radiology of the University Hospital in Leuven for their support.

## Funding

F.P. was funded by FWO (Fonds Wetenschappelijk Onderzoek) postdoc fellowship (12Q4615N) and research grant (1528216N). H.O.B. was supported by ERC-2011-Stg-284101 and IUAP P7/11. B.B. and H.O.B. were supported by Interdisciplinary Research Fund (IDO/10/003). H.O.B., F.P., B.B. were supported by FWO research grant (G088216N). J.B. was funded by FWO PhD fellowship (11J2115N). The funders had no role in study design, data collection and analysis, decision to publish, or preparation of the manuscript.

## Authors’ contribution

F.P., B.B. and HOB design; F.P., M.H., S.A., N.D. performed, F.P, J.B., H.L.M., HOB analyzed and F.P., J.B., B.B., H.O.B wrote the paper.

## Conflict of interest

The authors declare no competing financial interests.

